# Short-term heritable variation overwhelms two hundred generations of mutational variance for metabolic traits in *Caenorhabditis elegans*

**DOI:** 10.1101/2020.02.05.935197

**Authors:** Charles F. Baer, Dan Hahn, Lindsay M Johnson, Olivia J Smith

## Abstract

Metabolic disorders have a large heritable component, and have increased over the past few generations. Genome-wide association studies of metabolic traits typically find a substantial unexplained fraction of total heritability, suggesting an important role of spontaneous mutation. An alternative explanation is that epigenetic effects contribute significantly to the heritable variation. Here we report a study designed to quantify the cumulative effects of spontaneous mutation on adenosine metabolism in the nematode *Caenorhabditis elegans*, including both the activity and concentration of two metabolic enzymes and the standing pools of their associated metabolites. The only prior studies on the effects of mutation on metabolic enzyme activity, in *Drosophila melanogaster*, found that total enzyme activity presents a mutational target similar to that of morphological and life-history traits. However, those studies were not designed to account for short-term heritable effects. We find that the short-term heritable variance for most traits is of similar magnitude as the variance among MA lines. This result suggests that the potential heritable effects of epigenetic variation in metabolic disease warrant additional scrutiny.

## INTRODUCTION

Human metabolic diseases have increased markedly in frequency over the past few generations (Saklayen 2018). Large genome-wide association studies (GWAS) conducted on the human metabolome have shown that metabolic traits are highly heritable, but that a substantial fraction of the heritability of metabolic traits remains unexplained by the cumulative effects of mQTL (Rhee et al. 2013; Shin et al. 2014; Mahajan et al. 2018). This discrepancy indicates that the remainder of the heritable variation is the result of some combination of (1) rare, highly deleterious variants recently arisen in the population; (2) many variants with effects too small to be detected by the typical GWAS (Manolio et al. 2009; Eichler et al. 2010; Boyle et al. 2017); and/or (3) cross-generational epigenetic effects that are heritable over the short term but leave no genetic signature (Furrow et al. 2011; Richard et al. 2017). Scenarios (1) and (2) imply a significant role of spontaneous mutation in the risk of metabolic disease, although the rapid increase in frequency further implies some sort of genotype-environment interaction. A recent onslaught of epigenetic effects is considered less likely as a general explanation for the “missing heritability” of human complex traits (Wainschtein et al. 2019), but specific examples of cross-generational effects are known in humans (Pembrey et al. 2006; Curley et al. 2011; Veenendaal et al. 2013; Rando and Simmons 2015), and are well-documented in other organisms (e.g., plants; Munir et al. 2001; Luna et al. 2012; Rasmann et al. 2012) and *C. elegans*; (Greer et al. 2011; Rechavi et al. 2011; Ashe et al. 2012; Jobson et al. 2015; Marré et al. 2016).

To our knowledge, the cumulative effects of spontaneous mutation on metabolic traits have been investigated in only three experiments. Mukai et al. (1984) measured the cumulative effects of 300 generations of spontaneous mutations on the activity of alcohol dehydrogenase (Adh) in *Drosophila melanogaster*. In a groundbreaking study, also in *Drosophila melanogaster*, Clark et al. (1995) quantified the input of mutational (co)variance in the activity of a set of 12 metabolic enzymes and two metabolites. In both studies, mutational heritability (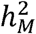, the per-generation increase in genetic variation (V_M_) scaled as a fraction of the residual variance, V_E_) of enzyme activity was on the order of that of life-history and morphological traits (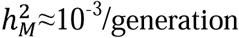; Houle et al. (1996)). In several of the mutation accumulation (MA) lines studied by Clark and his colleagues, there were large changes in enzymatic activity relative to the population mean over the course of 44 generations of evolution under minimal selection. Results for the two metabolites studied were analogous, but there was no attempt to assess the relationship between enzyme activity and metabolite concentration in the context of metabolic pathways.

More recently, Davies et al. (2016) examined the changes in metabolite concentration for 29 metabolites in a set of *C. elegans* MA lines that had undergone ∼250 generations of evolution under minimal selection and found that metabolites vary considerably in their response to spontaneous mutation, as quantified by the change in mean metabolite concentration (ΔM) and by the mutational (co)variance. Associations between mutational correlations between pairs of metabolites (*r*_*M*_, presumably the result of pleiotropy) and proximity of the metabolites in the global metabolic network were, on average, positive but weak (Johnson et al. 2018). The weakness of the association between mutational pleiotropy and network proximity suggests that pleiotropic effects propagate throughout the metabolic network and are not confined to local modules. However, there was no attempt to link changes in metabolite concentration to the properties of associated metabolic enzymes.

Here we report results of a study designed to investigate the cumulative effects of mutation on the concentration and activity of the enzymes in the adenosine metabolism pathway and their associated metabolites (Figure 1), using (nearly) the same set of *C. elegans* MA lines as in Davies et al. (2016). We chose this particular metabolic pathway for investigation because adenosine was one of the metabolites with the largest mutational variance, indicative of a large mutational target. In addition, adenosine levels are assumed to be tightly regulated due to its role as a critical signaling molecule for energetic homeostasis as a metabolite involved in ATP: AMP, as well has having other critical functions (Park and Gupta 2008; Boison 2013). Lastly, the adenosine pathway has well-defined network topology and is highly conserved.

**Figure 1.**
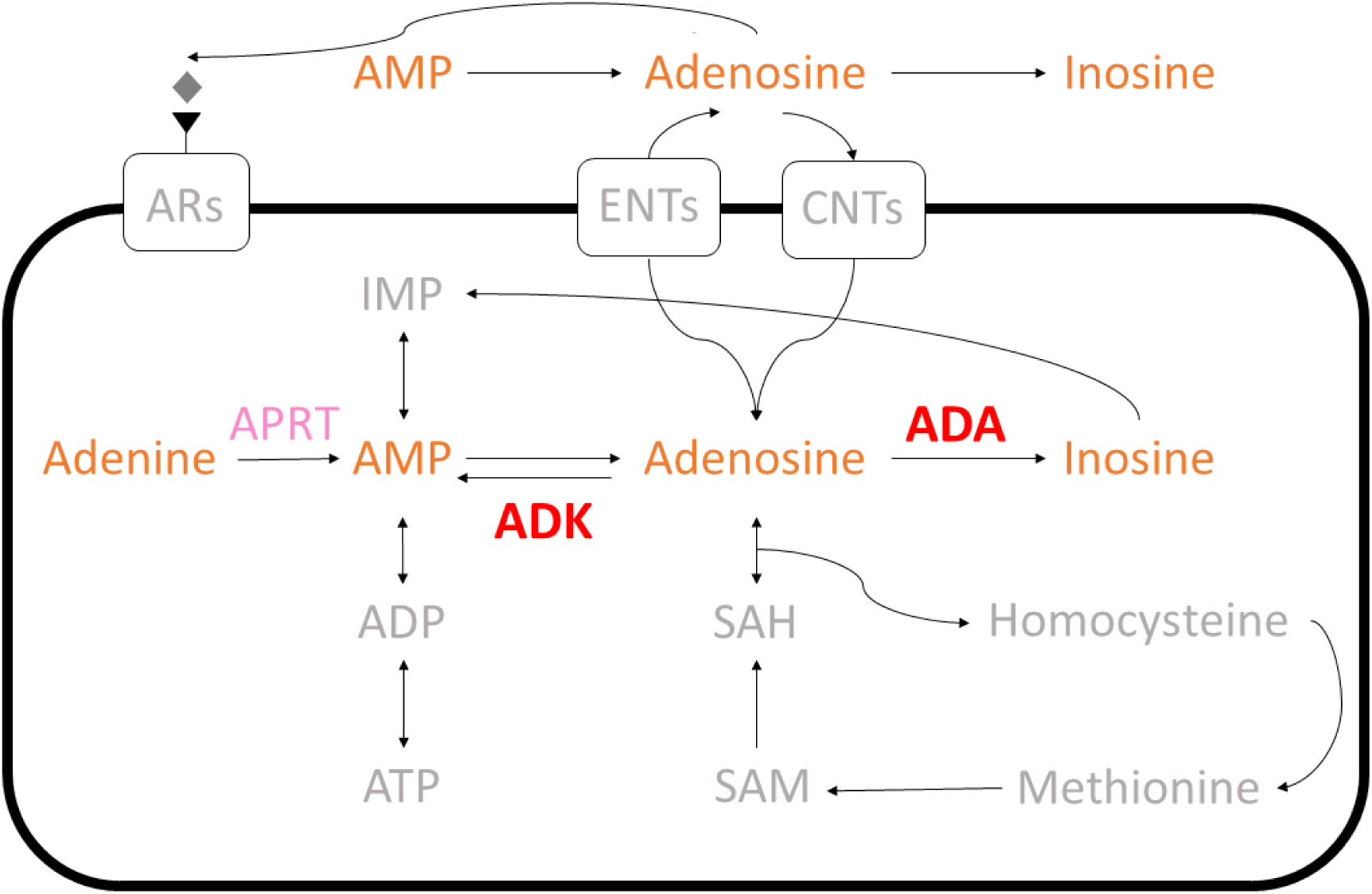
Adenosine metabolism pathway. Activity and concentration of enzymes Adenosine deaminase (ADA, red) and Adenosine kinase (ADK, red) were measured. We were unable to measure the concentration of APRT (pink). Metabolites in orange had concentrations quantified, those in gray were not measured.

Our study has one additional important feature relative to the aforementioned ones (Clark et al. 1995; Davies et al. 2016; Johnson et al. 2018). All of those studies estimate cumulative mutational parameters from the among-line components of (co)variance of a set of MA lines. Ideally, the among-line (co)variance is due solely to the contribution of new mutations, but other technical and biological factors can contribute to the among-line (co)variance (Lynch and Walsh 1998). Our experimental design allows us to infer the relative contributions of both mutation and short-term heritable (i.e., epigenetic) effects on the total heritable variance of metabolic traits.

## MATERIALS AND METHODS

### Mutation Accumulation

A detailed description of the construction and propagation of the mutation accumulation (MA) lines is given in Baer et al. (2005). Briefly, 100 replicate MA lines were initiated from a nearly isogenic population of N2-strain *C. elegans* and propagated by single-hermaphrodite descent at four-day (one generation) intervals for approximately 250 generations. The common ancestor of the MA lines (“G0”) was cryopreserved at the outset of the experiment; MA lines were cryopreserved upon completion of the MA phase of the experiment (Figure 2). Based on extensive whole-genome sequencing (Denver et al. 2012; Saxena et al. 2019), we estimate that the average MA line carries at least 60-100 mutant alleles in the homozygous state. In this study we included 39 of the 43 N2-strain MA lines assayed by Davies et al. (2016). One of the lines included in that study (line 507) was revealed by genome sequencing to be a contaminant from a different strain, and was removed from the analysis. Two other lines (517, 598) were revealed to have been cross-contaminated subsequent to the MA phase of the experiment, i.e., they appear to be genetically identical. Due to the structure of the experiment, we cannot simply pool the replicates of the two lines without introducing a potential bias, so those lines were omitted as well. All replicates of line 571 reproduced so slowly that we were unable to obtain sufficient material to be used in downstream assays.

**Figure 2.**
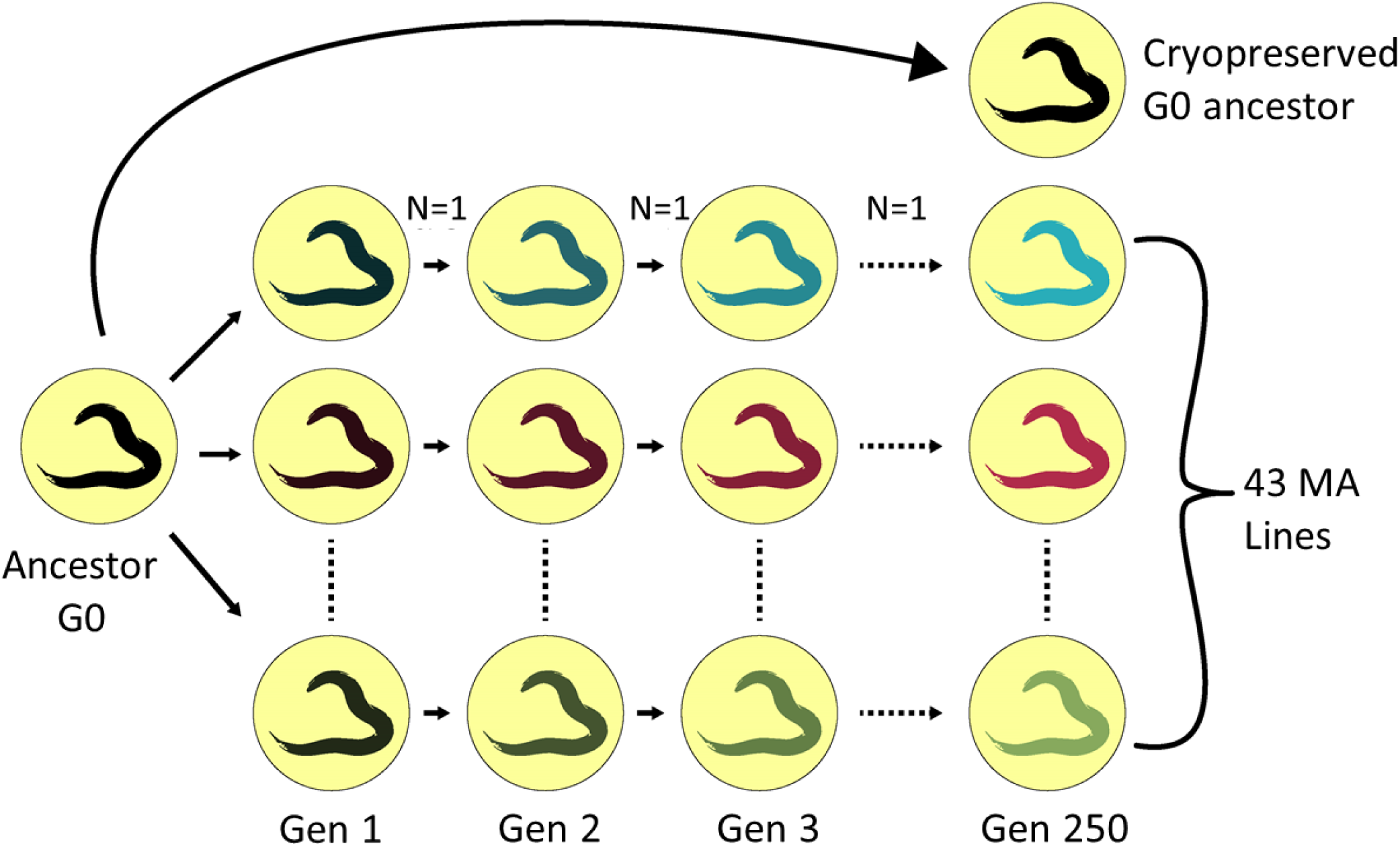
Propagation of mutation accumulation (MA) lines. The G0 ancestor was thawed from a cryopreserved sample and a single hermaphrodite picked onto each of 100 agar plates. MA lines were propagated via single worm descent for ∼250 generations. 43 MA lines and the G0 ancestor were included in this experiment.

The ideal design of a phenotypic assay of a MA experiment includes replicates of the (putatively) unmutated common ancestor, which we call “pseudolines” and which are treated identically to MA lines in analyses (Lynch 1985; Lynch and Walsh 1998; Teotónio et al. 2017). The among-pseudoline component of variance includes the effects of residual segregating genetic variation in the ancestor, as well as short-term heritable (epigenetic) effects that are propagated across assay generations and purely environmental effects resulting from (sometimes unavoidable) imperfections of experimental design, such as a temporal correlation between line and assay time. In the absence of a pseudoline control, some fraction of the among-MA line (co)variance will potentially be the result of non-mutational factors, and resulting estimates of V_M_ and COV_M_ will be upwardly biased.

Here, a set of 15 pseudolines (PS) of the G0 ancestor were included along with the MA lines (Figure 3A). PS lines were generated by thawing a sample of the N2 ancestor and allowing it 24 hours to recover from freezing, at which time 15 hermaphrodites were plated individually onto 60 mm NGM plates seeded with 100 *μ*l of an overnight culture of *E. coli* OP50 (P0 generation in Figure 3A). P0 worms were allowed to reproduce until the bacterial food on the plate was consumed (two generations; F1 and F2), at which time worms were cryopreserved (F2) (Hope 1999). The demographic features of this protocol mimic those of our standard protocol for cryopreserving MA lines. From this point forward, MA lines and ancestral PS lines are experimentally identical.

**Figure 3.**
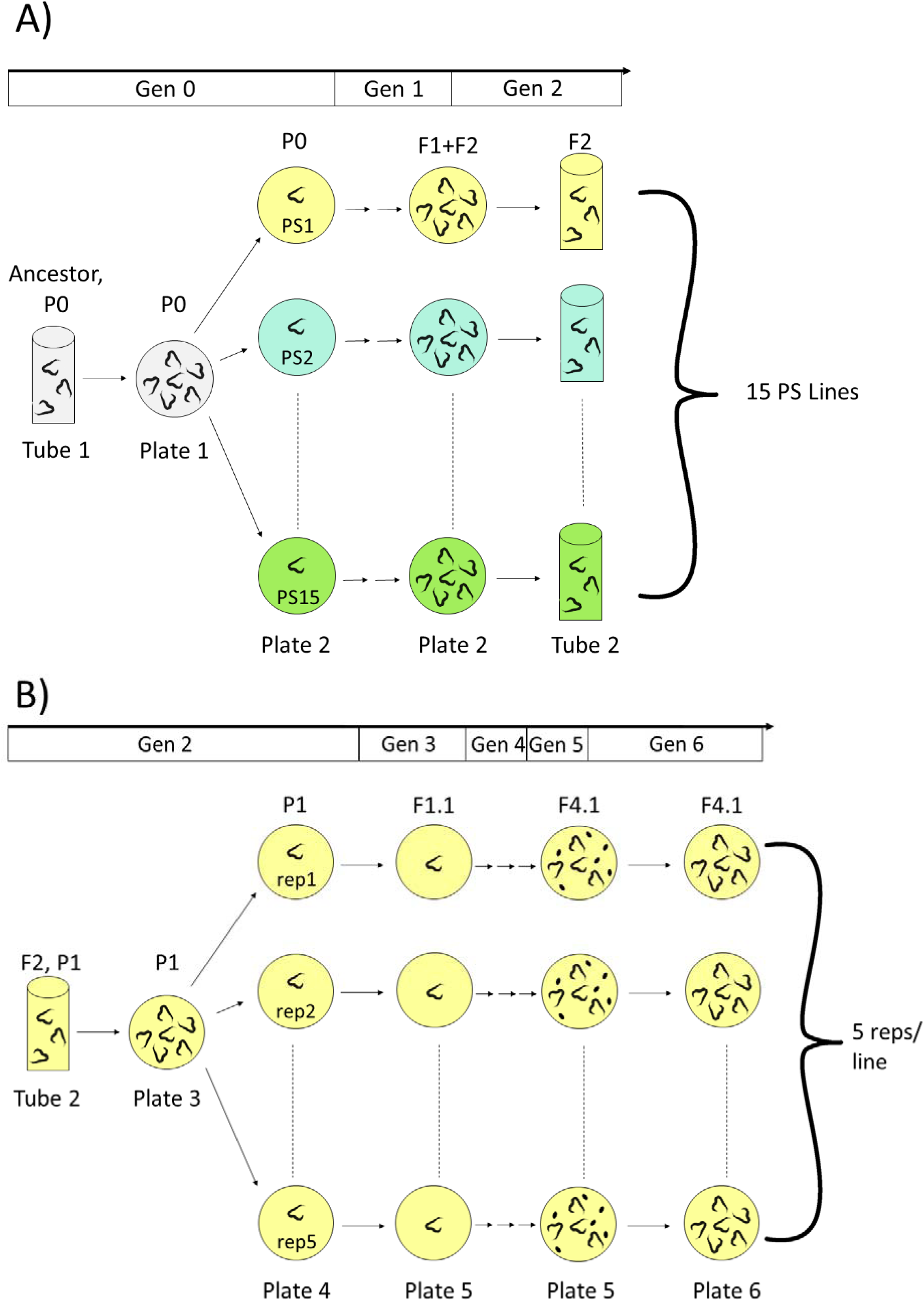
**A**) Generation of G0 pseudolines (PS lines). The G0 ancestor was thawed from a cryopreserved sample (“Tube 1”, “Plate 1”) and 15 individuals were picked onto individual agar plates (“Plate 2”; PS1-PS15) and allowed to reproduce for two generations prior to cryopreservation (“Tube 2”). **B**) Replication of lines for protein/metabolite extraction. Lines (P1, “Tube 2” from [A]) were thawed (plate 3) and five individuals were picked onto individual agar plates (“Plate 4”, Rep1-Rep5) and propagated by single-worm descent for another generation (F1.1, “Plate 5”). F1.1 worms were allowed to reproduce for two generations (F2.1, F3.1), and when the plates contained gravid worms (F3.1) and eggs (F4.1) they were bleached. The resulting eggs (F4.1) were transferred to a new plate (“Plate 6”) and allowed to hatch and grow to the young adult stage, at which time protein and metabolites were extracted. The timeline at the top represents the number of generations of reproduction of PS lines subsequent to divergence of the lines from the common ancestor. Population sizes at each generation are summarized in tabular form in Supplementary Table S1.

### Protein Extraction

This study includes six independent experimental tests: concentration and activity of two metabolic enzymes (ADA and ADK), total protein concentration, and mass spectrometry of pooled metabolites. We were unable to measure the activity of the third enzyme in the pathway, adenosine phosphoribosyltransferase (APRT), because commercially available assay kits require too much material to be practical for application to *C. elegans*. Accordingly, six aliquots of protein (plus metabolites) were extracted and cryopreserved from the same individual sample of each experimental replicate. Protein extraction was performed in five blocks of 10-12 lines per block, to ensure that all samples were handled at the appropriate stage of development (see below). In each protein extraction block, the lines selected were a random mix of MA and PS lines; the experimental design is outlined in Figure 3B. Each line was thawed and transferred onto a 60mm agar plate. The following day, five L4-stage hermaphrodites from each line were transferred individually onto 35mm agar plates (parental generation, P1 in Figure 3B), resulting in a total of 290 samples (five replicates of each of 15 PS lines and 43 MA lines). Four days later, a single offspring (F1 generation) L4 hermaphrodite was transferred from each P1 plate onto a 100mm plate (F1.1 in Figure 3B). The F1 worms were grown for ten days (two generations, F2.1 and F3.1 in Figure 3B) of self-replication to ensure that F3 worms were gravid and there were abundant eggs on the plate (F4.1 in Figure 3B). Worms were washed from the plate and “bleached” in an NaOH and sodium hypochlorite solution (Sulston and Hodgkin 1988). This process kills all hatched worms by breaking down their cuticle and leaves only eggs (F4.1 in Figure 3B), resulting in a population that is closely synchronized in developmental timing. Once F4 worms had been bleached, hatched, and reached the L4 stage, they were washed five times in ion-free NGM buffer, mixed with protease inhibitor cocktail, and homogenized via sonication (Tang and Choe 2015). Homogenized samples were centrifuged, and the protein-rich supernatant was distributed equally into six cryovials and stored at −80 C°. All lines, both MA and PS, were labeled with their true line number until cryopreservation, at which time each replicate was assigned a random number to obscure sample identity.

### Estimating Total Soluble Protein via Bicinchoninic Acid Assay (BCA)

We used total soluble protein as a proxy for the number of individual worms in a sample. To quantify the total soluble protein in each sample we used a bicinchoninic acid assay (BCA) following the protocol from Thermo Scientific (Pierce BCA Protein Assay Kit #23225). Briefly, a set of known concentrations of bovine serum albumin is used to generate a standard curve against which one can estimate the concentrations of unknown protein samples. A total of 13 BC assays were performed, each with its own set of standards.

### Enzyme activity assays

#### (i) Adenosine kinase (ADK)

Adenosine kinase (ADK) activity was measured using the Novocib PRECISE ADK assay kit (Novocib, Ref #K0507-01). This assay measures ADK activity based on the production on NADH_2,_ which is generated by the dephosphorylation of ATP by ADK. To ensure that ADK activity is not limited by available ATP, an excess of human ATP was added to each sample. Absorbance at 340nm was measured at one-minute intervals for 40 minutes. The slope of the line over the linear phase quantifies the activity of each sample in units of absorption per minute. A set of positive (human ADK, provided in the kit) and negative (no enzyme) controls were included with the unknown samples in each assay plate and used to quantify assay quality, per the manufacturer’s instructions. Thirty of the 290 samples were not included in the ADK activity assay because of erratic activity slopes. All samples that were run included at least two technical replicates, in which extracts from a sample were split and assayed independently.

#### (ii) Adenosine deaminase (ADA)

ADA activity was measured using Abcam’s Adenosine Deaminase (ADA) Activity Assay Kit (Abcam, ab21193). This kit utilizes an ADA developer and converter which react with inosine formed from the breakdown of adenosine by ADA to produce uric acid. Uric acid concentration is then measured via absorbance at 293nm once a minute for 45 minutes. Each kit is run with a set of known concentration standards that are used to generate a standard curve. The quantity of uric acid was then measured and used to calculate the activity of the ADA in a given sample in units of nmol/min/μg, following the manufacturer’s instructions.

ADA activity was assayed in six 96-well plates, each including a positive (manufacturer supplied ADA) and negative (no sample) control. For one assay plate, the highest concentration standard had an unusually low reading; we therefore omitted this point from the standard curve for this assay. Omission of that point had no effect on the interpretation of the data because all unknown samples had absorbance values greater than the second lowest standard. All of the 290 samples had maximum measured activity well below the highest concentration standard. Four samples with erratic absorption readings were omitted from further analyses.

### Enzyme concentration

Enzyme concentrations were estimated by Western blot (WB) (Supplemental Figure S1). Extracted samples were denatured in 2X Laemmli buffer (with β-mercaptoethanol) and boiled at 70° for 10 minutes. Each gel lane was loaded with 7ug of total soluble protein calculated from the BCA data (Bio-Rad 10% polyacrylamide gel, product #4561033). Each blot included eight samples, a DNA-ladder and an internal control standard consisting of a homogenate of *C. elegans*. We used the Trans-Blot Turbo Transfer System (Bio-Rad, #1704156) to transfer proteins separated by gel electrophoresis onto blotting paper. After the primary (enzyme-specific) and secondary (visualization) antibodies were bound (antibodies described below), antibody binding was visualized using the Pierce ECL Western Blotting Substrate (Thermo Fischer Product # 32106). Brightness of each band relative to the internal control was estimated using ImageJ image-analysis software and used as a proxy for enzyme concentration. 246 of the 284 samples contained sufficient protein to be visualized by Western Blot.

The concentration of tubulin in a sample is commonly used as a loading control, and we quantified tubulin in each sample for both enzymes (Tubulin antibody DSHB, E7). However, tubulin concentration was not independent of treatment (MA vs. PS), so we treat it as an experimental trait rather than a control (see Results).

#### (i) ADK concentration

The antibody used was Abcam’s Anti-ADK antibody – C-terminal (Abcam, ab226187), which was designed and tested in mouse and humans and which is homologous with the *C. elegans* ADK protein, R07H5.8. The assay resulted in multiple binding sites, with distinct bands at ∼100kd, ∼37kd, ∼25kd, and ∼18kd (Supplemental Figure S2). To determine which of these binding sites represented the *C. elegans* ADK, samples of each band were extracted from the gel and analyzed using protein mass spectroscopy. Results were then analyzed using Scaffold 4; only the sample at ∼37kd contained the worm ADK homolog (R07H5.8, molecular weight = 37.5 kd; Wormbase). 112 of the 246 samples did not contain sufficient ADK to be measured by Western blot. These lines were tested in duplicate and failed to produce ADK bands both times, therefore the low concentration of ADK is presumably a true property of the sample and not an experimental artifact.

#### (ii) Adenosine deaminase (ADA) concentration

The primary anti-body used was Abcam’s Anti-ADAT2 antibody (Abcam, ab122280). This antibody is homologous with the *C. elegans* ADA protein ADR-1 which is known to code for ADA in worms (Wormbase). The assay resulted in multiple binding sites, with distinct bands at ∼100kd, ∼60kd, and ∼22kd (Supplemental Figure S3). Samples of each band were extracted from the gel and analyzed using protein mass spectroscopy as for ADK. The band at ∼100kd contained the worm ADA homolog ADR-1, isoform D (101.8kd). 202 of the 246 samples contained ADA in sufficient concentration to be quantified by Western blotting.

### Metabolomics

To assess the relationship between enzyme concentration and activity and the concentration of their associated metabolites, we targeted four metabolites in the adenosine metabolic pathway: adenosine, inosine, AMP, and adenine. Several other metabolites not in the adenosine pathway were also measured, including GMP, guanine, guanosine, hypoxanthine, xanthine, and uric acid because they were part of a routine panel that included the metabolites of interest. Metabolite quantification was performed using liquid chromatography/mass spectroscopy (LC-MS), calibrated with known standards at the Southeast Center for Integrated Metabolomics at UF.

Internal standards were prepared as follows: Adenine-^15^N_2_ (Cat #A2880477), guanine-4,5-^13^C_2_ 7-^15^N (Cat #G836003), hypoxanthine-^13^C_2_ ^15^N (Cat #H998504) and xanthine-^13^C ^15^N_2_ (Cat #X499954) were purchased from Toronto Research Chemicals (Toronto, ON). Adenosine-^15^N_5_ 5′-monophosphate (Cat #662658), adenosine-^15^N_5_ 5′-triphosphate (Cat #707783), guanosine-^15^N_5_ 5′-monophosphate (Cat #900380) and guanosine-^13^C_10_ 5′-triphosphate (Cat #710687) were purchased from Sigma-Adrich (St. Louis, MO). The labeled adenosine and guanosine triphosphates were dephosphorylated with alkaline phosphatase (Promega, Madison, WI; Cat #M1821) according to the manufacturer’s directions to produce the corresponding labelled nucleosides. Uric acid-^13^C ^18^O was synthesized from urea-^13^C ^18^O (Cambridge Isotopes, Andover, MA; Cat #COLM-4861) and 5,6-diaminouracil sulfate (Sigma-Aldrich; Cat #D15103) according to methods of Cavalieri et al (Cavalieri et al. 1948).

For the purine assay, internal standard (10µl) was added to 50µl worm homogenate and acetonitrile (100µl) was added to precipitate proteins for LC-MS/MS analysis. Samples were chromatographed on a Waters Cortecs UPLC HILIC column (2.1 x 150 mm, 1.6µm) eluted with an acetonitrile-water gradient: Buffer A) 5 mmol/L ammonium acetate and 0.1% acetic acid in acetonitrile: water (:: 98: 2); Buffer B) 10 mmol/L ammonium formate and 0.5% formic acid in water. Mass spectrometric detection was on a Bruker EvoQ Elite MS/MS in positive ion mode, using heated electrospray ionization.

Stock solutions of the purines analyzed were prepared from authentic standards, and their concentrations determined by absorbance (Umbreit et al. 1960). The stock solutions were then mixed to give an appropriate working standard, which was then serially diluted to produce standard curves. Peak area ratios were calculated by dividing the metabolite peak area by the peak area of its isotopically labeled internal standard. Metabolite concentrations were calculated by comparing these peak area ratios to the standard curves.

### Data Analysis

#### (i) Estimation of mutational parameters

To quantify the cumulative effects of mutation on individual traits, we calculated the per-generation change in the average trait value (ΔM, the “mutational bias”) and the per-generation rate of increase in genetic variance (V_M_, the “mutational variance”). Mutational bias is calculated as: 

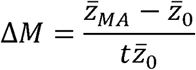

where 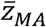 and 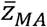 are the averages of the MA lines and the G0 pseudolines, respectively, and *t* is the number of generations of MA (*t*=250) (Lynch and Walsh 1998). We report values of ΔM using the median rather than the mean as the measure of the average because of the highly skewed distribution of some traits. Because protein and metabolite concentrations are not independent of the total amount of the sample, statistical tests of mutational bias are based on the slope of the linear regression of trait value on generations of MA (*R*_*M*_), including total protein concentration of the sample as a covariate.

The mutational variance (*V*_*M*_) is calculated as: 

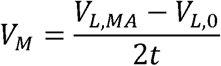

where *V*_*L,MA*_ is the among-line variance of the MA lines, *V*_*L,0*_ is the among-line variance of the PS lines, and *t* is the number of generations of MA. The among-line variance of the PS lines includes the effects of any residual segregating genetic variance, but also heritable epigenetic effects and the heritable effects of genotype-environment correlation (Lynch 1985).

The mutational covariance between traits (*COV*_*M*_) is estimated analogously to V_M_, with the among-line components of variance (*V*_*L*_) replaced with the among-line components of covariance (i.e., the off-diagonal elements in the variance-covariance matrix).

#### (ii) Statistical analyses

Our primary interest is in the two enzymes, ADA and ADK. The enzyme activity assays measure the composite effects of enzyme activity *per se* (i.e., the inherent kinetic properties of the protein) and the concentration of the enzyme in the sample. For a given sample, the rate at which substrate is converted to product depends on both the amount and the inherent activity of the enzyme present. Because we have an independent measure of the amount of enzyme present in the sample (from the Western blots), we can statistically partition the effects of inherent activity from those of concentration by including enzyme concentration as a covariate in a general linear model (GLM). The concentration of protein measured in the Western blot is standardized by the total protein in the sample, so enzyme activity also needs to be standardized relative to the total protein in the sample, which can be similarly included in a GLM. The ADA activity assay includes total protein in the calculation of activity, so total protein is not included in the GLM. The full GLM can be written as: 

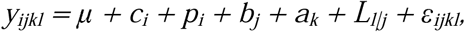

where *y*_*ijkl*_ is the measured activity of the enzyme in sample *i*, μ is the overall mean, *c*_*i*_ is the effect of the concentration of the enzyme in sample *i* (estimated from the Western blot of the same sample), *p*_*i*_ is the total protein concentration in sample *i, b*_*j*_ is the random effect of assay block *j, a*_*k*_ is the continuous fixed effect of generations of MA (PS=0, MA=250), *L*_*l|j*_ is the random effect of line *l* given MA group *k*, and ε*ijkl* is the residual effect. Variance components (V_L_ and V_E_) and their standard errors were estimated separately for each MA treatment group by restricted maximum likelihood (REML), with degrees of freedom determined by the Kenward-Roger method (Kenward and Roger 1997). Statistical significance of the change in trait means (*R*_*M*_) was assessed by F-test on Type III sums of squares, as implemented in the MIXED procedure of SAS v. 9.4.

Statistical significance of the among-line components of variance (V_L,MA_ and V_L,GO_) was assessed by randomization test, where trait values were randomized and variance components estimated from the GLM as previously described. Randomization was performed separately for MA and PS lines. Among-line variances (V_L,MA_ and V_L,G0_) were deemed significant if the point estimate was greater than 95% of 10,000 estimates from the randomized values. Randomization tests of variance components were performed using the lme4 package in R (Bates et al. 2015).

Protein concentrations (ADA, ADK, tubulin) were calculated relative to a predetermined amount of total protein (see section V above). Mutational statistics for protein concentrations were calculated from the same linear model as above without the covariates. Protein concentration data were log-transformed to meet the assumptions of the GLM, and statistical inferences are based on the transformed data. Mutational statistics are reported on the untransformed scale.

Metabolite concentrations were normalized relative to an internal standard. Mutational statistics were estimated from the same linear model as above, with total protein included as a covariate. Data were log-transformed to better fit the normality assumption of the GLM.

Because V_M_ is a composite measure (V_M_=V_L,MA_-V_L,PS_), in principle one could assess the statistical significance of an estimate of V_M_ by comparing the fit of a model with the among-line variance estimated separately for each treatment group (MA and PS) to that of a model with a single among-line variance component, e.g., by likelihood-ratio test. However, the difference in sample sizes between the two groups greatly reduces the power of the test. Instead, to assess the significance of estimates of V_M_, we employed a standard bootstrap approach (Baer et al. 2005), in which data are resampled with replacement at the level of line independently for MA and PS lines and variance components estimated as described above. 10,000 bootstrap estimates of V_M_ were generated for each trait; the upper and lower 2.5% of the distribution establish the empirical 95% confidence interval, which we use as our criterion of statistical significance.

Mutational (among-line) covariances were calculated in two ways. Our primary interest is in the correlations among the eight traits of the adenosine metabolic pathway. With eight variables, there are 8×9/2=36 elements in the half-diagonal covariance matrix, thus 39 MA lines are sufficient to jointly estimate the among-line components of covariance. The covariance was partitioned into within- and among-line components analogously to the GLM described above for variances, by REML with unstructured covariances as implemented in the MIXED procedure of SAS v. 9.4 (Fry 2004). For traits in which total protein is a relevant covariate, we first regressed the log-transformed value against the log of total protein concentration, then used the residual as the dependent variable in the GLM. For traits for which total protein is not a relevant covariate, we used the log-transformed value. Residuals of the full GLM were very close to normally-distributed, as assessed by visual comparison to a q-q plot. Statistical significance of among-line correlations was assessed by Wald’s Z-test.

For the full data set of 17 traits, there are 17×18/2=153 elements in the diagonal covariance matrix, too many to jointly estimate from 39 MA lines. Instead, we calculated pairwise correlations of line means, using the R package corr.test (Revella 2019). Trait values were standardized relative to the G0 mean across all PS lines.

## RESULTS

### Per-generation change in trait means (ΔM)

Box-plots showing line means and within-line variation are given in Supplementary Figure S4. Of the seventeen traits, only two (GMP and Xanthine) changed significantly over the course of 250 generations of MA at the experiment-wide 5% level of significance, although there was a general trend for metabolite concentrations to decline relative to the total protein in the sample (Table 1). Mean total protein concentration, by which the other trait values were standardized, was nearly identical in the G0 ancestor and in the MA lines. The close concordance in the average amount of total protein in a sample indicates that the average number of worms in a sample did not differ consistently between ancestor and MA lines. A caveat is in order, however. Although samples were synchronized by bleaching and were cultured to the same semi-objective stage of development (“a few” eggs were present on the plate), subtle differences in the distributions of developmental stages may exist at any hierarchical level in the experiment (G0 vs. MA; among lines; among replicates within a line). It is known that there are consistent changes in the genome-wide transcriptional profile over the course of a few hours of development (Francesconi and Lehner 2014; Zalts and Yanai 2017), and there is reason to expect that changes in metabolite levels would change at least as fast. Determining whether a given difference in trait value between two groups is due to a true difference in the trait at the exact same stage of development, or due to a subtle (perhaps non-linear) difference in rate of development is a nearly insoluble problem once worms have developed past a few embryonic cell divisions.

**Table 1.** Means. Column headings are: *M*_*MA*_, MA mean/median (SEM); *M*_*0*_, G0 pseudoline mean/median (SEM); *R*_*M*_, per-generation change in trait mean conditioned on total protein concentration (SEM); Δ*M*_*MED*_, per-generation change in trait mean scaled as a fraction of the G0 mean, calculated from median value of line means. Values of *R*_*M*_ in bold font are significantly different from zero (experiment-wide P<0.05). See Methods for details of the estimation of trait means.

| Trait | $M_{MA}$ | $M_0$ | $R_M$ (x $10^3$) | $\Delta M_{MED}$ (x $10^3$) |
| --- | --- | --- | --- | --- |
| ADA activity | 0.0037 / 0.0035<br>(0.00017) | 0.0038 / 0.0037<br>(0.00025) | -0.020 (0.414) | -0.216 |
| ADK activity | 0.012 / 0.012<br>(0.00005) | 0.012 / 0.012<br>(0.00014) | -0.020 (0.414) | 0.034 |
| ADA conc | 0.22 / 0.16<br>(0.034) | 0.41 / 0.29<br>(0.091) | -2.870 (1.266) | -1.746 |
| ADK conc | 2.71 / 0.71<br>(0.82) | 0.67 / 0.33<br>(0.20) | 2.004 (1.425) | 4.670 |
| Tubulin (ADA) | 0.18 / 0.17<br>(0.02) | 0.18 / 0.17<br>(0.020) | -1.010 (0.567) | -0.803 |
| Tubulin (ADK) | 0.18 / 0.18<br>(0.019) | 0.18 / 0.18<br>(0.019) | -0.900 (0.673) | -1.262 |
| Total protein | 0.70 / 0.66<br>(0.03) | 0.70 / 0.69<br>(0.037) | 0.111 (0.287) | -0.187 |
| AMP | 8.76 / 7.91<br>(0.74) | 11.26 / 9.75<br>(1.29) | -0.850 (0.717) | -0.756 |
| Adenine | 0.26 / 0.20<br>(0.026) | 0.28 / 0.20<br>(0.044) | 0.023 (0.762) | 0.012 |
| Adenosine | 1.75 / 0.35<br>(0.69) | 5.45 / 0.46<br>(2.50) | -1.870 (1.627) | -0.855 |
| GMP | 2.90 / 2.16<br>(0.28) | 4.36 / 3.76<br>(0.49) | **-2.380** (0.678) | -1.702 |
| Guanine | 1.42 / 1/31<br>(0.12) | 2.16 / 2.10<br>(0.33) | -1.360 (0.788) | -1.509 |
| Guanosine | 1.71 / 0.49<br>(0.55) | 4.00 / 1.04<br>(1.55) | -2.740 (1.553) | -2.106 |
| Hypoxanthine | 4.14 / 3.06<br>(0.56) | 5.21 / 5.20<br>(0.90) | -1.520 (0.825) | -1.647 |
| Inosine | 1.37 / 0.52<br>(0.34) | 2.52 / 1.04<br>(1.00) | -1.940 (1.156) | -1.998 |
| Uric Acid | 13.29 / 11/14<br>(1.20) | 17.30 / 12.05<br>(2.17) | -1.290 (0.568) | -0.299 |
| Xanthine | 3.03 / 2.34<br>(0.38) | 4.92 / 4.37<br>(0.59) | **-2.190** (0.651) | -1.864 |

As quantified in our enzyme activity assays, the output variable (“activity”) is a function of both enzyme kinetics (i.e., activity *per se*) and the amount of enzyme in the sample. Interestingly, for both enzymes, the correlation of the measured activity was either negatively or not correlated with the concentration of the enzyme. In the case of ADA, the negative correlation was highly significant (phenotypic correlation *r* = −0.34, df=176, P<0.001). The correlations for ADK were smaller and not statistically significant, albeit with a smaller sample size (*r* = −0.03, df=113, P>0.08). An obvious *post hoc* explanation is that the flux through the pathway is tightly regulated, and a change in activity *per se* is compensated for by a change in concentration of the enzyme in the appropriate direction, or possibly *vice versa*. That argument further implies, however, that the measured output of the reaction depends on factors other than the inherent activity of the enzyme itself, because at least in the PS lines the protein sequence is presumably identical in all samples (transcriptional and translational errors notwithstanding).

### Mutational variance (or the Lack Thereof)

As mutations accumulate over time, MA lines are expected to diverge in trait values, leading to a consistent, long-term increase in the among-line component of variance (V_L_). Scaled per-generation, this increase is the “mutational variance”, V_M_ (Lynch and Walsh 1998, p. 330). For various reasons, however, some fraction of the among-line variance may be due to factors other than the accumulation of new mutations. Possible reasons include residual segregating variation in the ancestor of the MA lines, genotype-environment correlations (sometimes unknown or unknowable), and heritable epigenetic effects (Rechavi and Lev 2017; Perez and Lehner 2019). To account for potential non-genetic contributions to the among-line variance, it is necessary to include a set of “pseudolines” (PS) of the ancestor, which are treated both experimentally and statistically as if they were MA lines (Lynch 1985; Teotónio et al. 2017).

We report two different standardizations of V_M_. First, the difference in the among-line variance between the PS and MA lines is divided by the square of the mean of the PS lines (V_M,0_); this is equivalent to the squared coefficient of variation, standardized by the ancestral mean. This quantity is often called the “evolvability” (Houle 1992), and is the customary way of scaling mutational variance. However, if the trait mean changes over the course of evolution, scaling the MA lines by the ancestral mean will underestimate the true mutational variance if mutational effects are multiplicative (i.e., the CV is constant; Fry and Heinsohn 2002; Baer et al. 2006). We also report V_M_ scaled by the group mean (V_M,MA_; i.e., PS lines are scaled by the square of the PS mean and MA lines are scaled by the square of the MA mean).

As it turns out, for most traits the among-line variance of the PS lines is of similar magnitude as that of the MA lines (Table 2; Supplementary Figure S4), with the result that none of the 17 traits showed significant mutational variance. Importantly, the lack of mutational variance is not because there is little among-line variance in the MA lines; in 14/17 cases V_L_ in the MA lines is significantly greater than zero (randomization test, p<0.05).

**Table 2.**
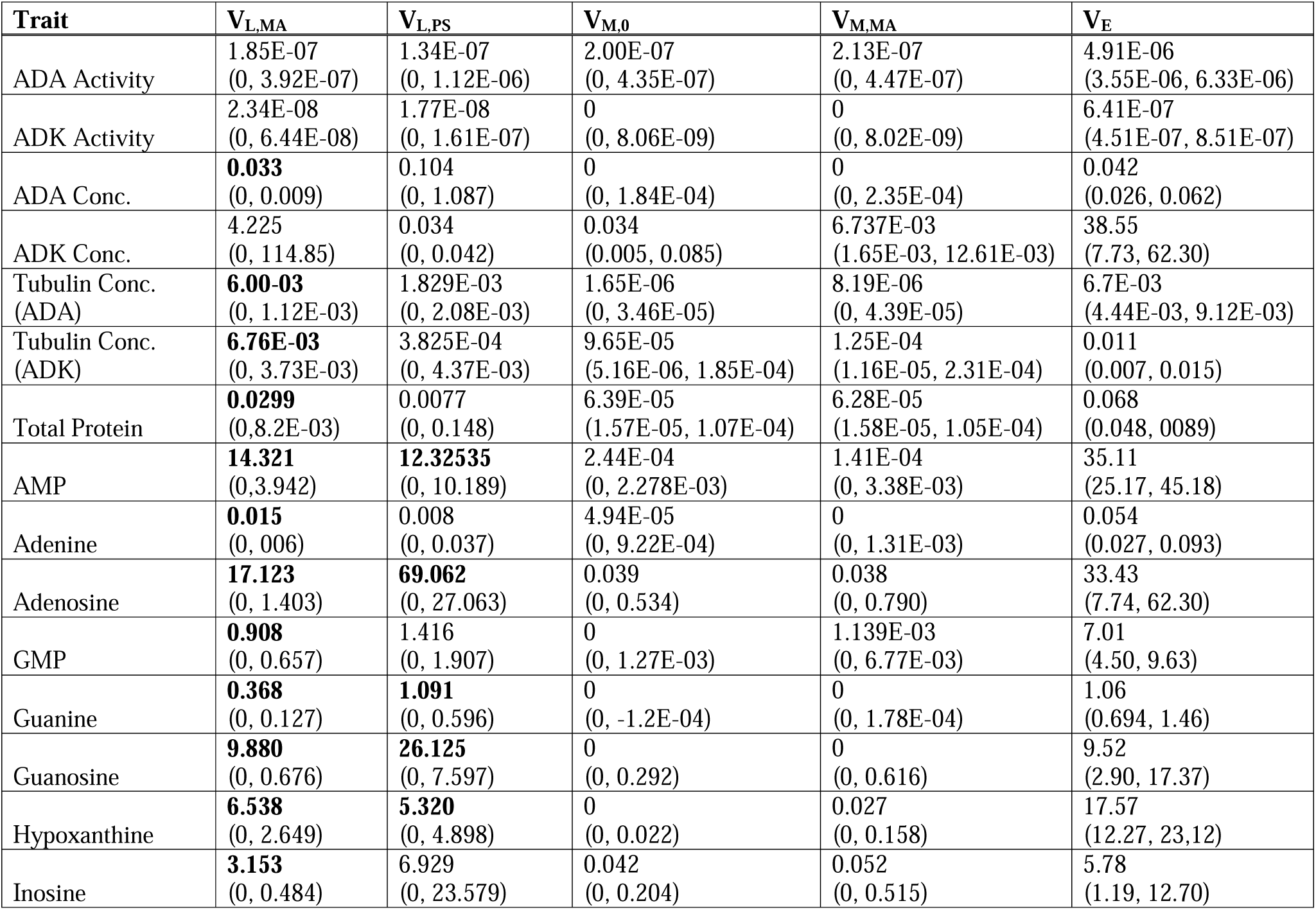
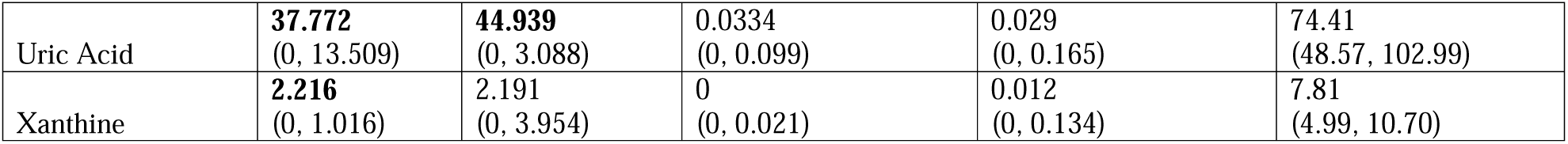
Variances. Column headings are: V_L,MA_, among-line variance of the MA lines; V_L,PS_, among-line variance of the G0 pseudolines; V_M,0_, the mutational variance standardized by the G0 mean; V_M,MA_, the mutational variance standardized by the mean of the group; V_E,_, the mean residual (within-line) variance of the MA and PS lines. Values of V_L_ and V_M_ in bold are significantly greater than zero; empirical 95% randomization confidence intervals are shown in parentheses for V_L,MA_ and V_L,PS_; 95% bootstrap confidence intervals are shown for values of V_M_. See Methods for details of the estimation of variance components.

Conceivably, technical variance associated with enzyme or metabolite assays could swamp biological variation and lead to a spurious partitioning of variance. However, several lines of evidence suggest this is not the cause of the substantial variance among PS lines. First, and most importantly, the technical variation would have to contribute in such a way as to inflate the among-line variance of the G0 pseudolines (i.e., a rather pathological Type 1 error), rather than inflating the among-biological replicate (within-line) variance and thereby simply reducing the power to detect among-line variance. Second, we ran technical replicates (i.e., samples of extracted material were split and assayed independently) for ADK activity. The among-technical replicate component of variance was about 1/3 that of the among-biological replicate variance, and pooling the technical replicates within a biological replicate or including them separately had no effect on the among-line variance. Based on previous experience with our metabolomics screen, technical replicate variance for the metabolic pools is expected to be less than 5% of the total variance for all metabolites except for GMP and uric acid, which are expected to be less than 10% (Eoin Quinlivan, Southeast Center for Integrative Metabolomics, personal communication).

It is also very unlikely that residual segregating genetic variance could explain the similar magnitudes of the among-line variance in the PS and MA lines. First, any residual genetic variation would be equivalently partitioned among PS lines and MA lines, and would contribute equally to the among-line variance (on average, sampling variance notwithstanding). The MA lines were initiated in March, 2001, at which time the G0 ancestor was expanded to large population size (three generations) and cryopreserved. Over the intervening 16 years prior to the start of this project, the ancestor has been thawed, re-expanded, and re-frozen several times. We do not know exactly how many times the ancestor has been thawed/expanded/re-frozen, but five is a reasonable guess. If we assume that each expansion takes three generations and there have been five such expansions, then any two PS lines will have diverged for 2×5×3=30 generations. In contrast, any two MA lines have diverged for 2x(250+3)≈500 generations. This is a conservative estimate because MA lines used for this experiment were also thawed and refrozen more than once, again we do not know how many times this has happened but it would inflate the number of generations of divergence between MA lines regardless of how many rounds of freeze-thaw occurred.

If technical and/or residual genetic variation cannot explain the among-line variance of PS lines, the most likely remaining possibility is heritable epigenetic effects. We cannot strictly rule out a vertically-transmitted pathogen, such as a virus or an intracellular parasite, e.g., microsporidia. However, there is no reason to expect variation in such a pathogen in long-term laboratory lines, whereas there is abundant evidence for heritable epigenetic effects in *C. elegans.* Recently, Sarkies and his colleagues reported that small RNA (specifically, piRNA/22G RNA) epimutations accumulate spontaneously at a rate ∼25X that of DNA sequence mutations, with a half-life on the order of 2-3 generations, but with significant fraction maintained for ten generations or more (Beltran et al. 2019). That time-scale is entirely consistent with the findings reported here. Protein and metabolite data were collected on the F4 descendants of the most recent common ancestor of a line (Figure 3B; Supplemental Table S1), which means that any non-genetic short-term heritable effects that are common to a line had to have been maintained for at least four generations, and perhaps since the founder of the PS line six generations back (Figure 3A). Thus, effects common to a line meet the definition of “transgenerational” effects (i.e. passed down to at least the F3, Rechavi and Lev 2017). We return to the topic of epigenetic inheritance in the Discussion.

### Among-line correlations

The absence of significant mutational variance precludes estimation of mutational covariances, which was one of the underlying motivations of this study. However, because there is significant among-line variance for most traits in both the PS and MA lines, it is meaningful to investigate the among-line correlations (Figure 4; Supplementary Figure S5). Note that these are not phenotypic correlations in the usual sense. Presumably, the among-line correlations reflect what might be thought of as epi-pleiotropy – the effects of an epigenetic variant (whatever it may be) on multiple traits – as well as the cumulative pleiotropic effects of new mutations.

**Figure 4.**
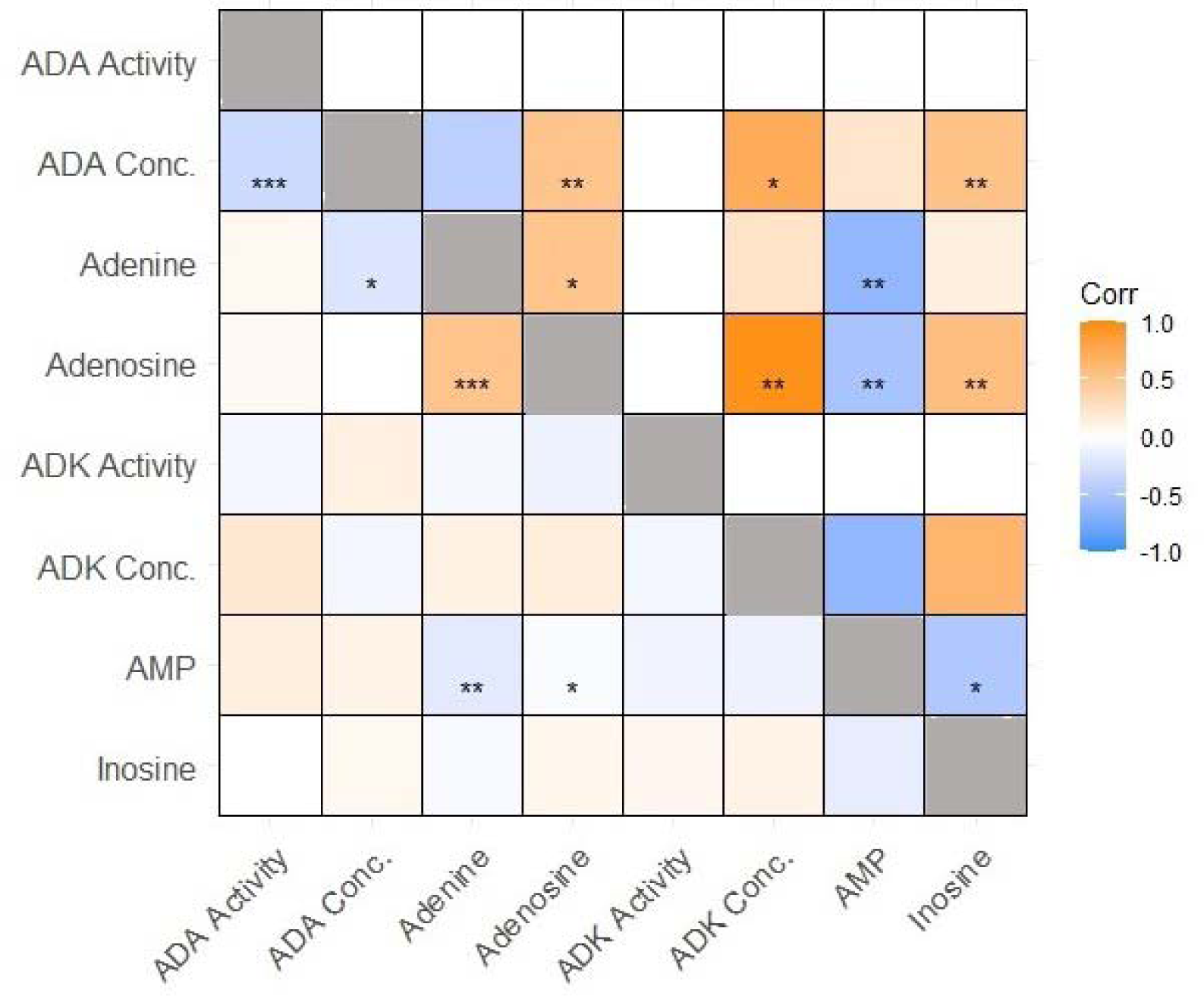
Heatmap of Pearson’s correlations between traits within the adenosine pathway. Among-line correlations are shown above the diagonal, within-line correlations (V_E_) are shown below the diagonal. Wald’s test significance levels are represented by: *** = p<0.001, ** = p<0.01, *=p<0.05.

There were significant positive among-line correlations between the concentration of ADA and its substrate (adenosine; (*r* = 0.51, p<0.01) and product (inosine; *r*= 0.53, p<0.01), and between the pools of adenosine and inosine (*r* = 0.56, p<0.01). The among-line correlation of ADK concentration with that of its substrate adenosine was large and positive (*r* = 0.96, p<0.01), whereas the correlation of the concentration of ADK with its substrate AMP is negative, but estimated with large variance (*r* = −0.63, p>0.09). Pools of AMP and adenosine were negatively correlated (*r* = −0.52, p<0.01). We were unable to measure the activity or concentration of adenosine phosphoribosyltransferase (APRT), which converts Adenine to AMP. Looked at more broadly, the absolute values of the point-estimates of 25 of the 28 among-line correlations within the adenosine metabolic network were greater than those of the same within-line correlation, irrespective of statistical significance. The among-line and within-line correlations were of the same sign in 19/28 cases, and in none of the nine cases in which the correlations were of opposite sign was the among-line correlation significantly different from zero.

## DISCUSSION

We find ourselves confronted with an inconvenient truth: taken at face value, the results we report here are starkly contradictory to the findings reported in Davies et al. (2016). We chose the adenosine metabolism pathway for further scrutiny based on two findings of Davies et al. In that study, mean adenosine concentration *increased* by over 4% per generation – one of the largest values of ΔM reported for any trait in any organism – whereas in this study we found a (non-significant) decline in adenosine concentration of about 0.1%/generation in (nearly) the same set of MA lines (Table 1). Similarly, Davies et al. reported a mutational heritability (V_M_/V_E_) for adenosine concentration of about 0.004/generation – toward the high end of mutational heritabilities (Houle et al. 1996) – whereas, we found no significant mutational variance. The discrepancy is not restricted to a handful of traits: Davies et al. reported significant mutational variance for 22 of the 29 metabolites included in their study. Clearly, the two studies are at odds: they can’t both be right, although they may both be wrong in different ways. The discrepancy is not due to the exclusion of three lines from this study, which had little effect on the results for most traits in our data (Supplementary Figure S4). The methods of quantifying metabolite concentration were different in the two studies; we used LC-MS in this study, whereas Davies et al. used GC-MS, but a poor workman blames his tools.

Critically, the lack of mutational variance is not because there is no variation between MA lines. For every trait except ADK activity and concentration, the variance among MA lines (V_L,MA_) is significantly greater than zero (Table 2). The cumulative effects of mutation are not swamped by technical or microenvironmental noise (i.e., residual variance; V_E_ in the parlance of quantitative genetics). Rather, the variance among pseudolines of the ancestral control is of similar magnitude to the variance among MA lines. Had we made the common assumption that there was no heritable variance in the G0 ancestor, our results would have been entirely consistent with those reported by Davies et al. (and the majority of MA studies reported in the literature, e.g., Clark et al. 1995).

For economic reasons (metabolomics is expensive), Davies et al. did not include pseudolines of the G0 ancestor in their study. Returning to adenosine as an exemplar, all but three of the 43 MA lines included in the Davies et al. study had mean adenosine concentrations greater than that of the G0 ancestor, which was an order of magnitude less than the mean of the MA lines in normalized units (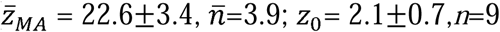; see Figure 1 of Davies et al. (2016)). Because ΔM is measured relative to the ancestor, if the mean value of the ancestor is atypically small, ΔM will be atypically large. We have no reason to doubt the accuracy of the estimate of mean adenosine concentration of the G0 ancestor in the Davies et al. study. 3/43 MA lines had mean concentrations lower than the ancestor, and another seven MA lines had means less than the largest of the nine replicates of the ancestor. Moreover, the average metabolite concentration of the ancestor was not low relative to the MA lines when all 29 metabolites are considered: the median rank of the ancestor is 34/44 (data from Davies et al. (2016) are archived in Dryad, at https://datadryad.org/stash/dataset/doi:10.5061/dryad.2dn09).

It is important to carefully consider the differences between the ways the ancestral controls were treated in the two studies. At the outset of the Davies et al. study, in 2009, a single cryopreserved sample of the ancestor was thawed in the Baer lab (Florida) and plated. From that plate, a “chunk” containing hundreds of worms was transferred onto another plate and sent to the Leroi lab in England, at which time worms were washed from the plate and cryopreserved at - 80° C. Later, one tube of the ancestor was thawed and plated onto a 100 mm plate. When the population on that plate reached high density (2-3 generations), worms were washed from the plate and “bleached” (Sulston and Hodgkin 1988), and surviving L1 larvae were chunked onto a new plate. From that plate, nine replicate plates were initiated from a single individual, and the populations grown to high density (2-3 generations) and synchronized by bleaching. Surviving L1s were plated and grown until worms reached young adulthood, at which time worms were collected for extraction of metabolites. In this design, the nine replicate plates are conceptually identical to the five replicates of each MA line, and the among-replicate (=within-line) variance is the residual variance, V_E_.

In this study (depicted in Figure 3A), 15 replicate plates were initiated from a single individual, grown to high density (two generations), and cryopreserved. These are the 15 ancestral pseudolines (PS). Subsequent to thawing (depicted in Figure 3B), the PS lines were treated identically to MA lines, with five replicate plates per PS line initiated from a single individual worm taken from the thawed plate. The replicates were then propagated to the F3 descendants of the original founder of the replicate, and their offspring (F4) collected for analysis. The variation among replicates is the residual variance, V_E_. Any effects that are common to a PS line (i.e., which contribute to V_L_) must necessarily have been maintained at least since the replicates diverged from their most recent common ancestor four generations previously, and potentially for as many as the six generations subsequent to the founding of the PS lines.

We believe the source of the discrepancy in ΔM between the two studies is likely the same as the source of the discrepancy in V_M_: short-term heritable, epigenetic variation. For example, there is a ∼120X difference between the mean adenosine concentrations between the two most extreme of the 43 MA lines in the Davies et al. study. The conventional interpretation is (and was) that spontaneous mutations accumulated over a couple of hundred generations can lead to huge differences in metabolite concentrations (and presumably in the concentrations of other biological molecules). However, there is a ∼100X difference in the mean adenosine concentration between the two most extreme of the PS lines in this study, lines that have diverged for only a few generations. If the one aliquot of the ancestor sampled in the Davies et al. study just happened by chance to fall in the lower tail of the distribution, voilà: ΔM “among the largest reported for any trait” (quoting Davies et al. 2016, p. 2243).

Given that the short-term heritability observed here is in fact epigenetic, what might be the cause(s), both proximate (i.e., mechanistic) and ultimate (e.g., environmental)? There is a burgeoning literature on heritable epigenetic effects in *C. elegans*, which can have a number of mechanistic causes, including several varieties of small RNA (Rechavi and Lev 2017), histone modifications (Furuhashi et al. 2010; Rechtsteiner et al. 2010; Tabuchi et al. 2018), and possibly 6-methyl adenine in DNA (Greer et al. 2015). Heritable epigenetic effects have been shown to affect a wide variety of traits (Schott et al. 2014; Demoinet et al. 2017; Han et al. 2017; Kishimoto et al. 2017), and in some cases have been shown to last for tens of generations (Ashe et al. 2012; Rechavi and Lev 2017). Parental age (Perez et al. 2017) and nutrition status (Miersch and Doring 2012; Tauffenberger and Parker 2014; Jobson et al. 2015) are especially well-documented drivers of epigenetic variation and are obvious potential sources of variation in the experiments reported here.

Nailing down the mechanistic cause(s) responsible for the epigenetic variation inferred here would be both very interesting and very challenging, but it is beyond the scope of this study. The most promising avenue of investigation would seem to be an experiment in which samples were split for combined metabolomics/transcriptomics, with a focus on piRNA/22G RNA variation (Beltran et al. 2019). However, while we do not know the mechanistic underpinning(s) of the apparent epigenetic variation among the ancestral pseudolines, the fact that we detected so much variation suggests that it is an important consideration in mutation accumulation studies, and more generally, in any quantitative genetic study in which phenotypic variance is partitioned within and among genotypes. Whether the high short-term heritability applies to taxa other than worms is unknown. However, a study of DNA-methylation in a set of *Arabidopsis thaliana* MA lines revealed that 5-methyl-cytosine epimutations occurred at a frequency several orders of magnitude greater than base substitution mutations (Becker et al. 2011). The dominant modes of epigenetic control differ between plants and nematodes (*C. elegans* apparently does not methylate cytosine in DNA), but the general conclusion that epimutations can introduce potentially important heritable effects in the short term is unavoidable.

In the only study comparable to this one, Clark et al. (1995, Table 3) found significant mutational heritability for the activity of 8/12 metabolic enzymes in a set of ∼50 *Drosophila melanogaster* MA lines that had evolved under MA conditions for 44 generations. However, their assay conflates variation in enzyme activity *per se* and variation in enzyme concentration into the composite category “enzyme activity” (normalized by body weight and total protein concentration), without correcting for enzyme concentration. The *Drosophila melanogaster* genomic mutation rate is perhaps 3X greater than that of *C. elegans* (Sharp and Agrawal 2012; Schrider et al. 2013), which suggests that after 44 generations of MA, a Drosophila MA line would have accumulated approximately half as many mutations as one of our *C. elegans* MA lines. Contrary to our expectation based on the preceding evidence, neither of the two metabolic enzymes we assayed (ADA and ADK) exhibited among-line variance for activity *per se* in either the MA lines or the PS lines. Thus, for those traits, we cannot attribute the absence of V_M_ to the confounding effects of among-line variance in the ancestor. It is interesting that the activity of these two enzymes is similarly unperturbed by both mutation and epigenetic factors. However, neither ADA nor ADK was included in the CLARK et al. study; it is certainly possible that had those enzymes been included in that study, they would have fallen in the group of enzymes without significant V_M_.

We conclude with two thoughts. First, and more parochially, for this set of metabolic traits (enzyme activity notwithstanding), a few generations of short-term heritable (presumably) epigenetic effects swamp the signal of ∼250 generations of accumulated mutations. Perhaps that should not be surprising: it is simply phenotypic plasticity, albeit of a different sort than evolutionary biologists are used to thinking about. It does strongly suggest, however, that investigators doing MA experiments need to be especially mindful of how the ancestor is treated, or employ designs in which direct comparison to an ancestor is not needed, such as regression of the among-line variance on generations of MA over multiple assays at different time points. In fact, that was the design employed in the Clark et al. study, but they constrained the intercept to equal zero, on the assumption that the among-line variance of the ancestor was zero. But also, second, and more broadly: these findings cast the recent increase in human metabolic complex disease in a different light. Although we remain skeptical of epigenetic variation as a general cause of “missing heritability” in humans, it may be that metabolic traits are particularly susceptible to epigenetic regulation and are worthy of closer scrutiny in that regard.

## Supporting information

Raw data and readme file

