## Supplemental figures and tables for "Short-term heritable variation overwhelms two hundred generations of mutational variance for metabolic traits in *Caenorhabditis elegans*"

| **Gen label** | **Gen count** | **Tube/plate** | **N worms** | | **N lines** | | **N reps/line** |
| --- | --- | --- | --- | --- | --- | --- | --- |
| P0 | 0 | tube1 to plate1 | Many | 1 | | 1 | |
|  | - | plate1 to plate 2 | 1 | 15 | | 1 | |
| F1 | 1 | plate2 | 200 | 15 | | 1 | |
| F2 | 2 | plate2->tube2 | 200^2 | 15 | | 1 | |
| F2->P1 | - | tube2 to plate3 | many | 15 | | 1 | |
| P1 | - | plate3 to plate4 | 1 | 15 | | 5 | |
| F1.1 | 3 | plate4 to plate5 | 1 | 15 | | 5 | |
| F2.1 | 4 | plate5 | 200 | 15 | | 5 | |
| F3.1 | 5 | plate5 | 200^2 | 15 | | 5 | |
| F4.1 | 6 | plate5 | many eggs | 15 | | 5 | |
| F4.1 | - | plate5 bleach to plate6 to extract | many | 15 | | 5 | |

**Table S1:** Pseudoline (PS) propagation and replication information.

| **Trait** | **Mean MA** | **Mean PS** | **∆M * 1000** |
| --- | --- | --- | --- |
| ADA Activity | 0.004 | 0.004 | -0.008 |
| ADA Conc | 0.205 | 0.782 | -23.060 |
| Adenine | 0.324 | 0.159 | 6.576 |
| Adenosine | 1.350 | -2.465 | 152.613 |
| ADK Activity | 0.012 | 0.012 | 0.002 |
| ADK Conc | 2.802 | 0.419 | 95.339 |
| AMP | 8.165 | 14.024 | -234.339 |
| GMP | 0.557 | 3.088 | -101.265 |
| Guanine | 1.031 | 1.631 | -24.007 |
| Guanosine | 0.632 | -0.086 | 28.711 |
| Hypoxanthine | 1.195 | -2.375 | 142.810 |
| Inosine | 0.177 | 5.635 | -218.331 |
| Total Protein | 0.700 | 0.697 | 0.143 |
| Tub ADA | 0.130 | 0.167 | -1.466 |
| Tub ADK | 0.135 | 0.180 | -1.765 |
| Uric Acid | 11.022 | -0.311 | 453.324 |
| Xanthine | 0.521 | -1.203 | 68.975 |

**Table S2:** MA and G0 means calculated from raw data via lmer described in methods. ∆M is calculated as MA mean minus the PS mean, divided by the number of generations of MA (250).

**A**

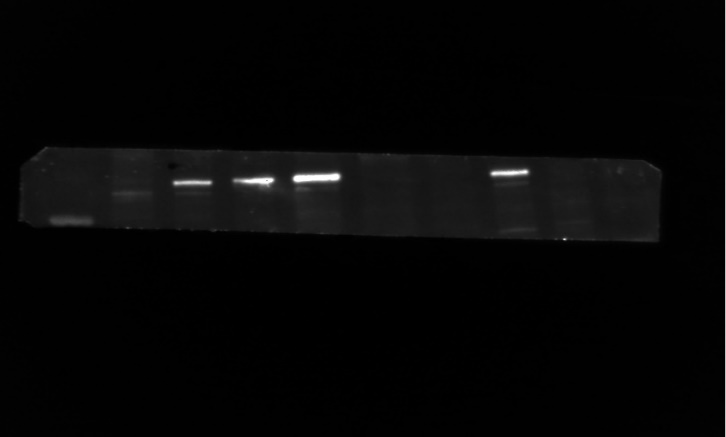

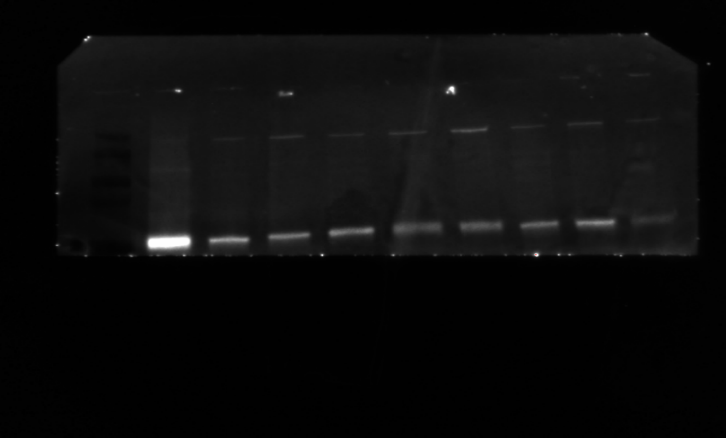

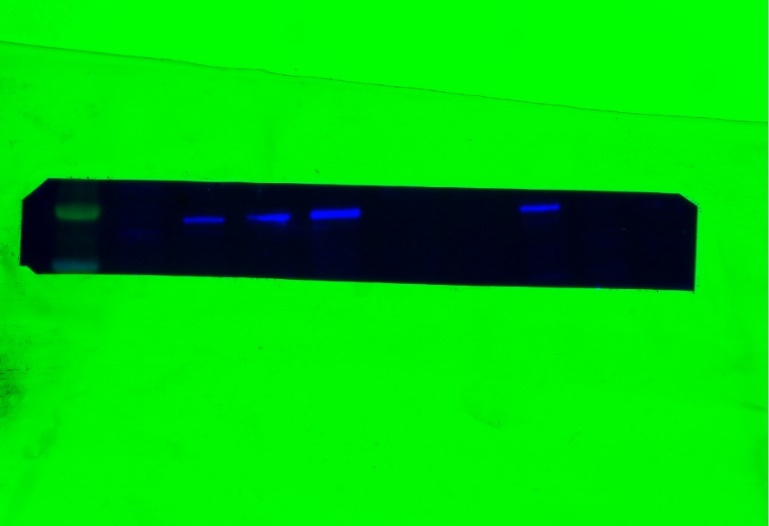

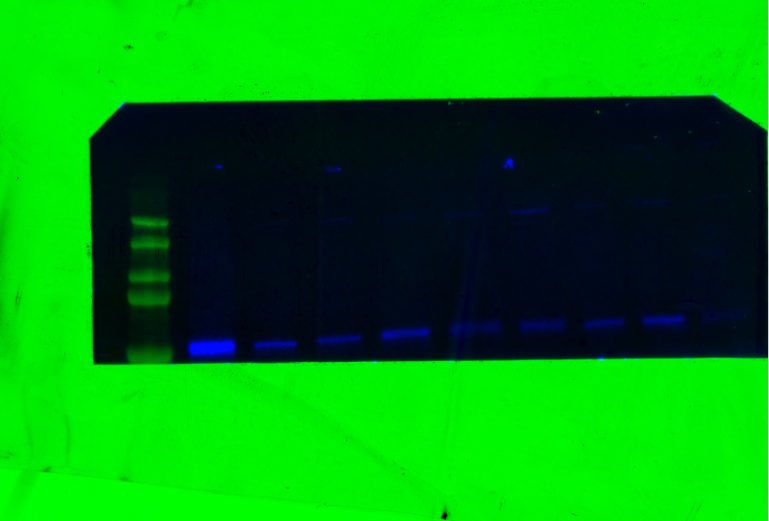

**B**

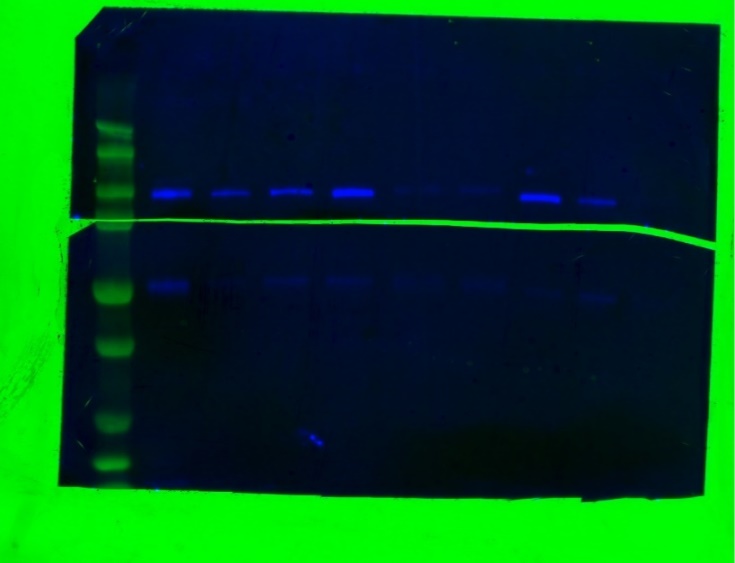

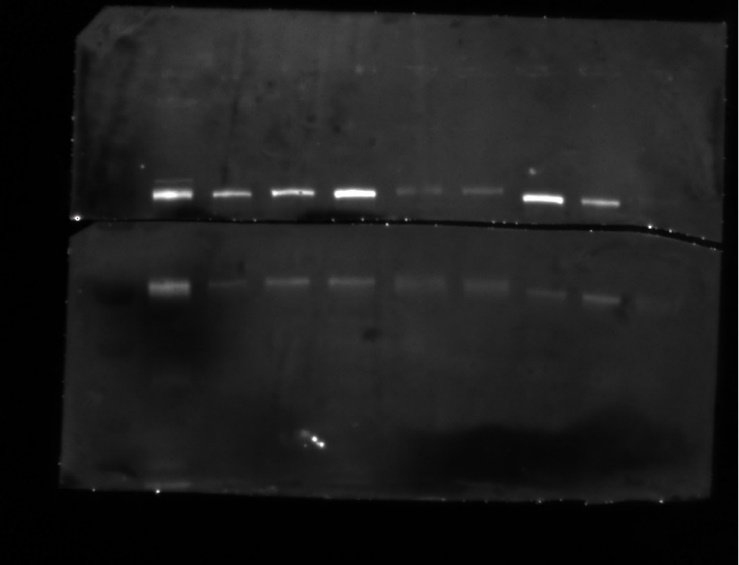

**Figure S1:** A) Adenosine kinase WB (color and black and white), top bands represent tubulin binding (red box), lower bands represent ADK binding (purple box). Lane 1 contains weighted DNA ladder, lanes 2-10 are samples 61-70. B) Adenosine deaminase WB (color and black and white), top bands represent ADA antibody binding (yellow box), lower bands represent tubulin binding (red box). Lane 1 contains weighted DNA ladder, lanes 2-10 are samples 61-70.

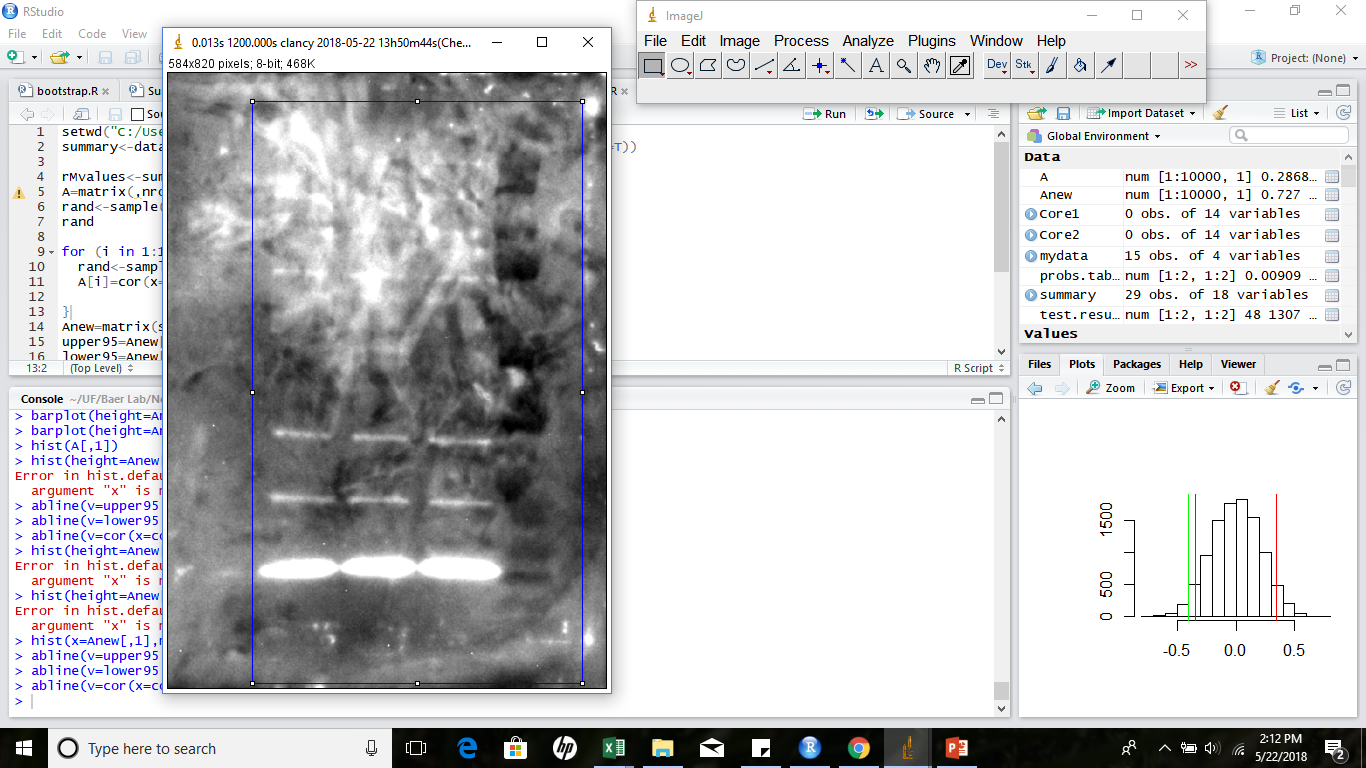

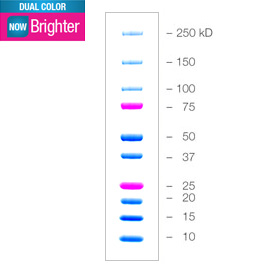

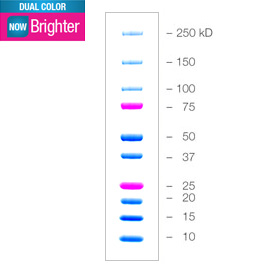

**Figure S2:** Adenosine kinase western blot with non-specific binding at ~200kb, ~37kb, ~25kb, and ~18kb. Lades from left to right contain: 1) 10ul worm protein denatured with 10ul β-mercaptoethanol denatured at 70° for 10 minutes; 2) 12ul worm protein denatured with 10ul β-mercaptoethanol denatured at 70° for 10 minutes; 3) 15ul worm protein denatured with 10ul β-mercaptoethanol denatured at 70° for 10 minutes; 4) 15ul DNA ladder.

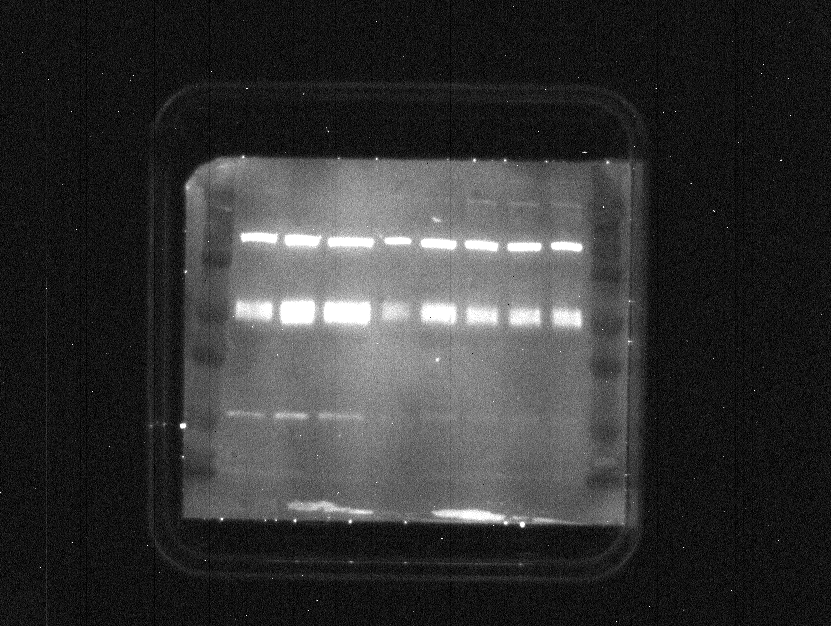

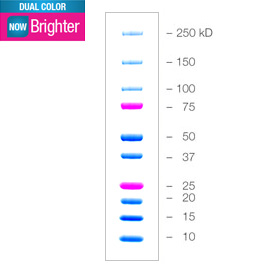

**Figure S3:** Adenosine deaminase WB with non-specific binding at ~100kb, ~60kb, and ~22kb. Lane contents from left to right: 1) 15ul DNA ladder; 2) 10ug worm protein with 10ul β-mercaptoethanol denatured at 70° for 10 minutes; 3) 10ul worm protein with 10ul β-mercaptoethanol denatured at 95° for 10 minutes; 4) 10ul worm protein with 10ul β-mercaptoethanol denatured at 95° for 5 minutes; 5) 10ul worm protein with 10ul DDT denatured at 95° for 5 minutes; 6) 10ul worm protein with 10ul DDT denatured at 95° for 10 minutes; 7) 10ul worm protein with 10ul DDT denatured at 70° for 10 minutes; 8) 10ul worm protein with 10ul DDT denatured at 50° for 20 minutes; 9) 10ul worm protein with 10ul DDT denatured at 50° for 10 minutes; 10) 15ul DNA ladder.

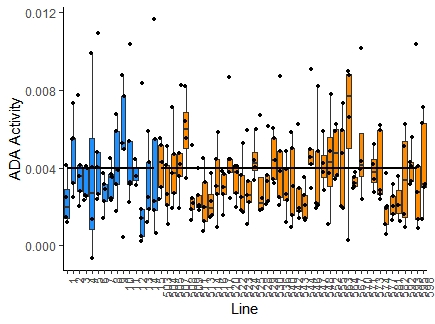

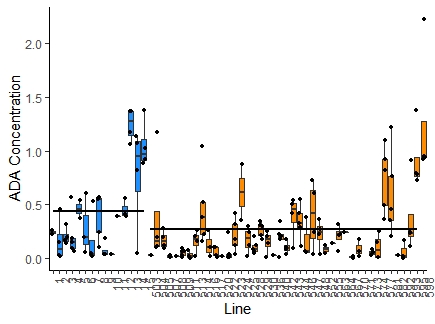

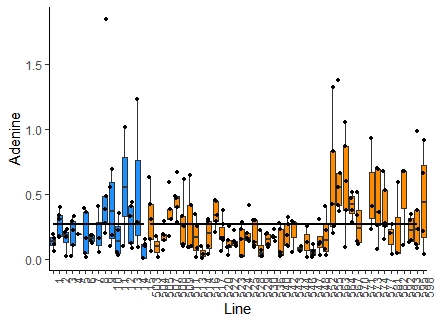

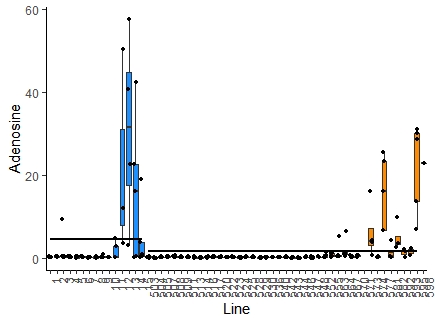

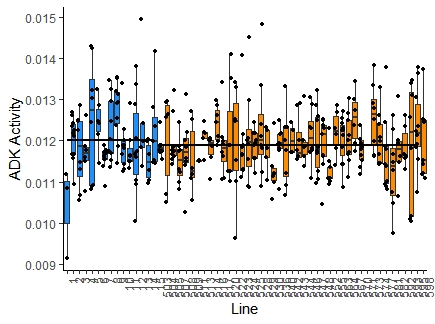

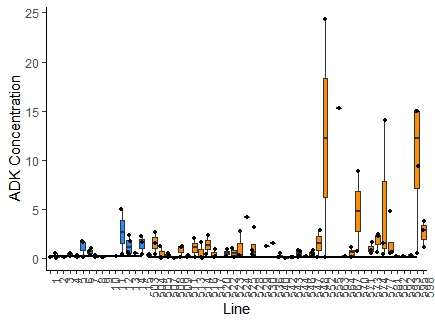

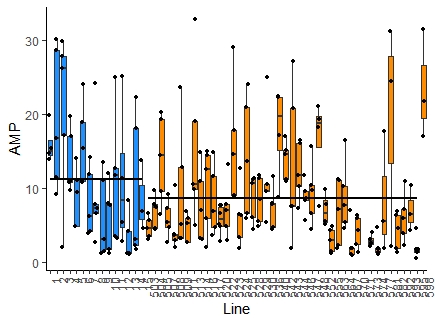
.
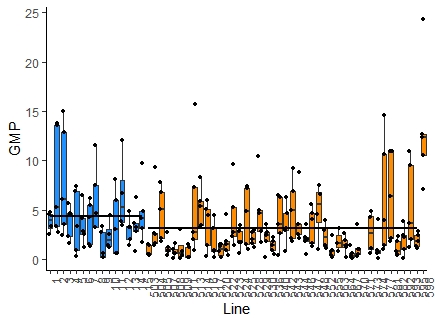

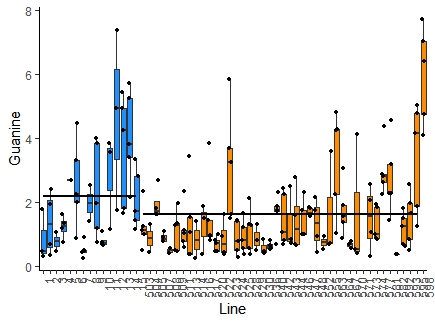

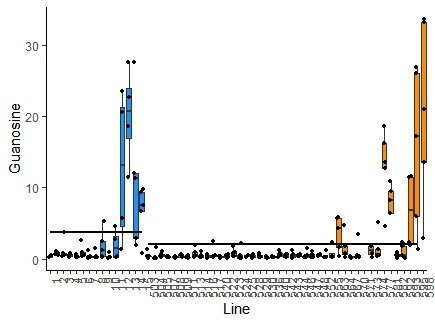

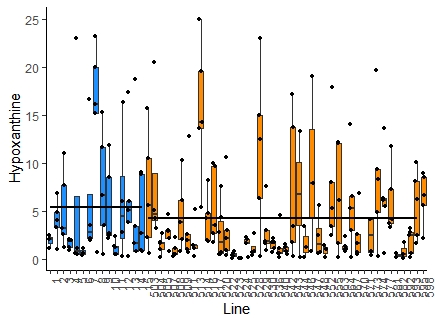

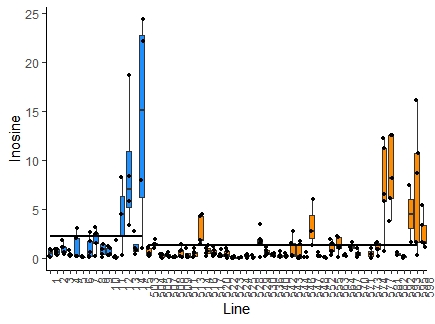

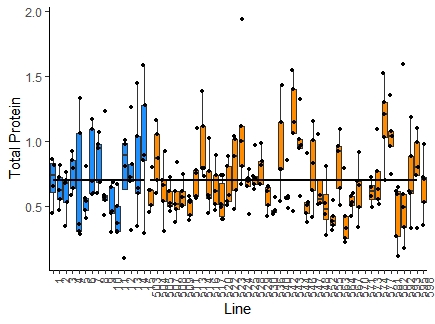

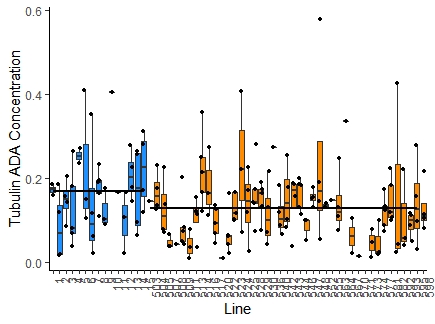

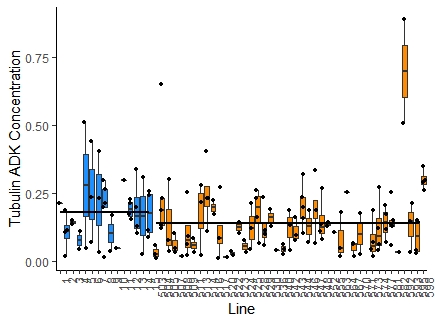

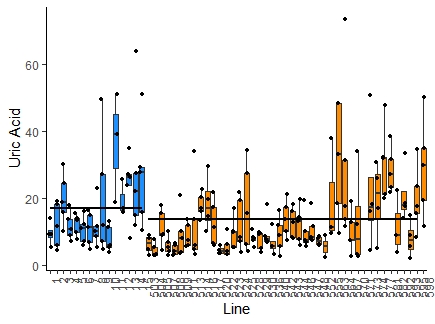

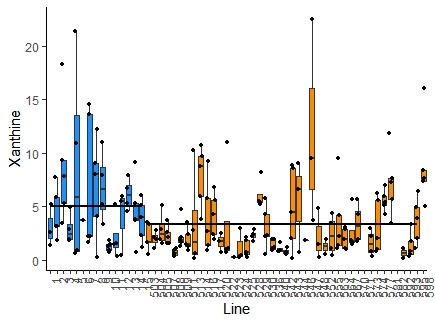

**Figure S4:** Box plots of trait values, by line; black dots are values for individual replicates. Trait identity is shown on the y-axis and lines are arranged in numerical order on the x-axis. G0 pseudolines (PS) lines are depicted in blue and MA lines are shown in orange. Black horizontal lines represent mean trait values of MA and PS lines respectively. MA line 507 (fourth MA line from left) is a contaminant and is not derived from the N2 strain. MA lines 517 (tenth MA line from left) and 598 (rightmost line) are genetically identical. Lines 507, 517, and 598 were removed from the analyses.

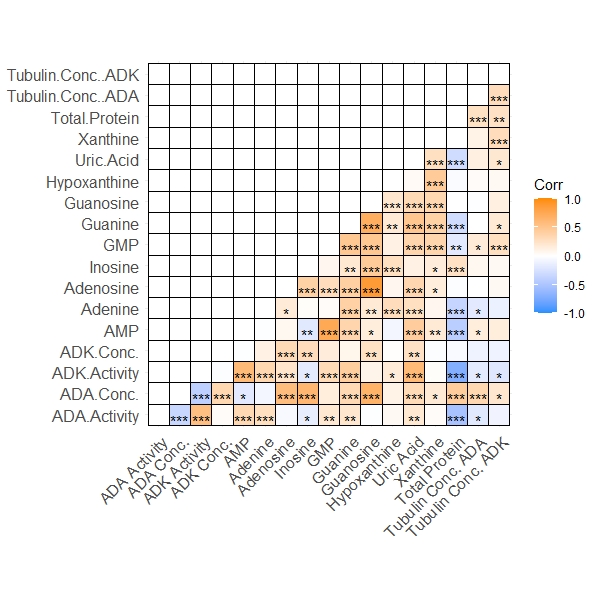

**Figure S5:** Heatmap of correlations of line means for all traits. Significance levels are shown as follows: *** = p<0.001, ** = p<0.01, *=p<0.05.
